# Quantifying bacterial evolution in the wild: a birthday problem for *Campylobacter* lineages

**DOI:** 10.1101/2020.12.02.407999

**Authors:** Jessica K. Calland, Ben Pascoe, Sion C. Bayliss, Evangelos Mourkas, Elvire Berthenet, Harry A. Thorpe, Matthew D. Hitchings, Edward J. Feil, Jukka Corander, Martin J. Blaser, Daniel Falush, Samuel K. Sheppard

## Abstract

Measuring molecular evolution in bacteria typically requires estimation of the rate at which nucleotide changes accumulate in strains sampled at different times that share a common ancestor. This approach has been useful for dating ecological and evolutionary events that coincide with the emergence of important lineages, such as outbreak strains and obligate human pathogens. However, in multi-host (niche) transmission scenarios, where the pathogen is essentially an opportunistic environmental organism, sampling is often sporadic and rarely reflects the overall population, particularly when concentrated on clinical isolates. This means that approaches that assume recent common ancestry are not applicable. Here we present a new approach to estimate the molecular clock rate in *Campylobacter* that draws on the popular probability conundrum known as the ‘birthday problem’. Using large genomic datasets and comparative genomic approaches, we use isolate pairs that share recent common ancestry to estimate the rate of nucleotide change for the population. Identifying synonymous and non-synonymous nucleotide changes, both within and outside of recombined regions of the genome, we quantify clock-like diversification to estimate synonymous rates of nucleotide change for the common pathogenic bacteria *Campylobacter coli* (2.4 x 10^-6^ s/s/y) and *Campylobacter jejuni* (3.4 x 10^-6^ s/s/y). Finally, using estimated total rates of nucleotide change, we infer the number of effective lineages within the sample time-frame – analogous to a shared birthdays – and assess the rate of turnover of lineages in our sample set over short evolutionary timescales. This provides a generalizable approach to calibrating rates in populations of environmental bacteria and shows that multiple lineages are maintained, implying that large-scale clonal sweeps may take hundreds of years or more in these species.

**Author Summary:** Growth and reproduction in living organisms require DNA replication but this process is error prone. Along with variation introduced by horizontal gene transfer, it can lead to alterations in the nucleotide sequence. These nucleotide changes accumulate over time in successive generations at an approximately constant rate termed the molecular clock. Therefore, if this rate is known, one can estimate the date when two or more lineages diverged. In bacteria, this can be informative for understanding the time-scale of emergence and spread of pathogenic strains. Such analyses are robust when the ancestral population is known, such as for obligate pathogens that only infect humans. However, when the bacterium inhabits multiple hosts or niches it is difficult to infer direct ancestry from one strain to another, reducing the accuracy of molecular clock estimates. Here we focus on one such multi-host organism, *Campylobacter*, a leading cause of food-borne gastroenteritis. Reconstructing the population history by estimating empirical nucleotide change rates from carefully selected isolate pairs, and evaluating the maintenance of multiple lineages over time, we provide information about strain diversification. Our method is a new addition to the bacterial genomics toolkit that will help in understanding the spread of opportunistic pathogens.

## Introduction

Theoretical models of a relatively constant rate of molecular change over time [1], the molecular clock, have become fundamental to explaining the evolution in bacteria [2, 3]. Spurred by the increasing availability of population-scale genome datasets, it is now common for comparative genomic studies to describe not only the relatedness of isolates but also how long ago they diverged [4–8]. This can provide valuable information when combined with host, habitat or ecosystem data. For example, it is possible to investigate how events such as host transitions or global dissemination have influenced the emergence and spread of lineages that may display important phenotypes, including pathogenicity.

There are significant challenges when applying molecular clocks to date lineage diversification in natural bacterial populations. In particular, it is necessary to determine the rate at which the clock ‘ticks’ and the uniform accumulation of nucleotide change (NC) over time. However, this is not simply a reflection of the background point mutation rate (associated with replication error) and the generation time of the bacterium [9, 10], but is also influenced by horizontal gene transfer (HGT) that can introduce several NCs in a single event [11]. Furthermore, the rate at which NCs accumulate in the population is influenced by the population size [12] and selection (positive and stabilizing) on different fitness effects [13].

While debate continues about NCs that are effectively neutral, and hence provide accurate clock estimates [14], there is clear utility for even approximations of the rate of genome change over time [15, 16]. This has allowed the development of time-calibrated phylogenies explaining molecular evolution in numerous well-known pathogen species [4–7]. However, even with large genome datasets and increasingly sophisticated models [17, 18], the accuracy of molecular evolution estimates is dependent upon the data from which they are derived, and two important considerations remain. First, the data should represent a longitudinal sample set [15, 19]. Second, the data should be representative of the population as a whole.

It is conceptually simple to understand how a long time-frame between collection of the earliest and latest sample would increase the number of NCs recorded, and how sampling at consistent intervals could help to determine if accumulation was linear over time. Comparisons between modern samples and DNA from the stomach of a 5,300 year old frozen iceman ‘Otzi’ have been used to investigate the emergence of modern *Helicobacter pylori* lineages [20]. However, ancient pathogen samples are rarely available. More frequently, molecular clock rates are estimated using collections of contemporary isolates that often share a common ancestor older than the sample frame. Convincing estimations have been possible for medically important bacteria, through comparison of large numbers of closely related isolates [21–24] but for many pathogens sampling of outbreaks may not provide an adequate representation of the bacterial population.

Most disease-causing bacteria are not obligate human pathogens. In this case, large reservoirs of isolates from which infection can arise may be infrequently sampled, despite their potential importance as emergent pathogenic strains. For example, *Campylobacter jejuni* and *C. coli* are among the most common causes of bacterial gastroenteritis worldwide but exist principally as commensal organisms in the gut of mammals and birds [25–29]. Human infection results primarily via food contaminated with strains from wild and agricultural animals, especially chickens [30–35]. In multi-host (niche) transmission scenarios such as this, where the pathogen is essentially an environmental organism, sampling is often sporadic and rarely reflects the overall population, particularly when concentrated on clinical isolates [36].

Overcoming the problem of sporadic or unrepresentative sampling for molecular clock estimation requires that sufficient numbers of isolates are collected to ensure that there are pairs that share a recent common ancestor (within the sampling period). However, with the enormous effective population size of environmental bacteria populations, questions remain about how many isolates need to be sampled to achieve this. This is analogous to the well-known probability theory conundrum known as the birthday problem [37]. This puzzle asks how many randomly chosen people need to be sampled so that a pair of them will share the same birthday. To be sure, requires a sample size of 366 (the number of possible birthdays), assuming that all birthdays are equally common, but a 99.9% probability is achieved with just 70 people and 50% with 23 people. This may seem counter intuitive but can be explained by considering that rather than comparing the birthday of a single individual to everyone else’s, in fact comparisons are made between every pair of individuals, 23 x 22/2 = 253. The result is greater than half the number days in the year, hence the 50% probability. Clearly, there are challenges in relating this conceptual model to bacteria. First, it is not known how many possible lineages (here equivalent to birthdays) there are in natural bacterial populations. Second, how to define lineages or isolate pairs with recent common ancestry. Third, just as with birthdays, some lineages are far more common than others. For example, of >72,000 *C. jejuni* and *C. coli* isolates archived in the pubMLST database [38], >50% belong to just 5 clonal complexes (out of 45).

Together, factors relating to isolate sampling and genome analysis conspire such that it may be difficult to distinguish NCs that reflect the passage of time [16, 39]. Here, we take a multi-layered approach to estimate the rate of molecular evolution of *C. jejuni* and *C. coli* using a large genome collection (2,425 genomes) representing isolates sampled over a 46-year period. We begin by identifying closely related isolate pairs in which the most recently sampled isolate has accumulated NCs over time. We then quantify synonymous and non-synonymous polymorphisms to take (some) account of selection, both within and outside of recombinant regions of the genome, and use synonymous polymorphisms to quantify clock-like diversification in *Campylobacter* [40, 41]. Finally, using estimated rates of nucleotide change we assess the rate of turnover of lineages in our sample sets over short evolutionary timescales. This provides a generalizable approach to calibrating rates in populations of environmental bacteria and clues about lineage diversification in two important enteric pathogens.

## Results

### There is a weak temporal signal in *C. coli* and *C. jejuni* phylogenies

Core genome phylogenies revealed little evidence of clustering by collection date (Fig 1). Isolates belonging to common sequence types (STs) and clonal complexes were sampled over the 46-year period. For *C. coli* and *C. jejuni* respectively, 1, 16, 3, 211, 370 and 41, 3, 34, 469, 1277 isolates were sampled over 50, 40, 30, 20, and 10 years ago. These included poultry associated ST-353, ST-354 and ST-257 complexes, cattle associated ST-61 and ST-42 complexes, and host generalist ST-21, ST-45, ST-828 (*C. coli*) complexes [42] (Fig 1 **and S1 Table**). Linear regression of root-to-tip distances and sampling dates of *C. coli* and *C. jejuni* phylogenies **(S1 and S2 Fig)**, using TempEst software, provided very weak evidence of a temporal signal when the best-fitting root was estimated. The R^2^ values were low for both *C. coli* (R^2^ = 0.176, slope = 3 x 10^-5^) and *C. jejuni* (R^2^ = 9.5 x 10^-2^, slope = 6.4 x 10^-5^) phylogenies (**S3 and S4 Tables**). Root-to-tip regression analysis was also run for *C. coli* and *C. jejuni* on three separate phylogenetic trees which were built from core gene alignments with the top 1, 5 and 10% most and least variable core genes (alleles / locus) filtered and removed. However, temporal signal remained poor for both *C. coli* (Avg. R^2^ = 2.4 x 10^-2^) and *C. jejuni* (Avg. R^2^ = 1.3 x 10^-2^) (**S3 Fig**). Consistent with some other studies [43], this poor branch-length to isolation date correlation suggests that estimation of the molecular clock rate from the entire dataset may be difficult. However, the accumulation of polymorphisms exhibited a positive correlation with sampling date in all datasets (**S1 and S2 Figs**) implying the maintenance of multiple STs and clonal complexes through time.

**Fig 1.**
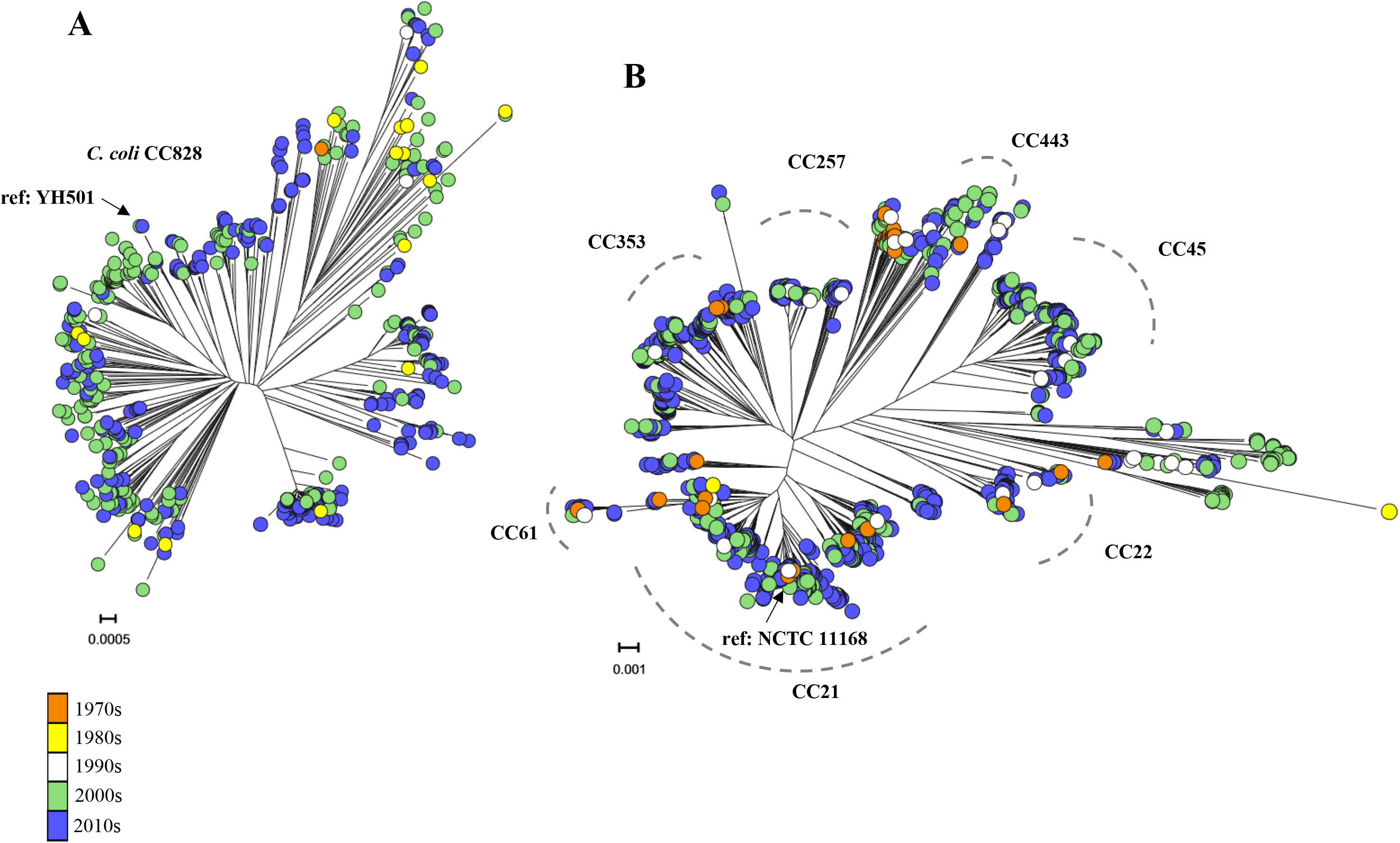
Little evidence of clustering of isolate sampling dates in *Campylobacter* phylogenies. Maximum likelihood (ML) core genome phylogenetic trees of *C. coli* (A) (n = 601) and *C. jejuni* (B) (n = 1824) constructed using FastTree version 2.1.8 [79] and the *GTR* model of nucleotide evolution. Both phylogenies show the distribution of the sample time frame used in this study with major *Campylobacter* clonal complexes (CCs) identified and terminal nodes coloured according to isolation decade (orange = 1970s, yellow = 1980s, white = 1990s, green = 2000s, blue = 2010s). Scale bars represent the estimated number of NCs per site. Terminal nodes sampled from different decades can be seen scattered throughout both trees with little evidence of clustering by decade. Isolates sampled from the 2000s and 2010s are most abundant within each dataset. The position of the *C. jejuni* (NCTC11168) and *C. coli* (YH501) reference genomes are indicated on the phylogeny. These were sampled in 1977 and 2016 respectively.

Analyses of temporal signal can be improved in bacterial genomes by the removal of recombined regions where multiple genetic variations may be introduced in a single evolutionary event [44]. Masking recombination in this way is challenging for large genome datasets such as those used in this study. Therefore, we conducted root-to-tip regression analysis on sub-lineages (tree clusters) within *C. coli* and *C. jejuni* where recombination events could be efficiently excluded (**S3 and S4 Tables**). Consistent with previous studies [44], any lineage with an R^2^ value >0.5 was described as having a strong temporal signal and was used in subsequent Bayesian evolutionary analysis. There were large differences in the strength of the temporal signal across all sub-lineages with the lowest R^2^ value found in the host generalist *C. jejuni* ST-45 clonal complex (R^2^ = 0.0229, slope = 1.89 x 10^-5^) and highest in *C. coli* ST-1090 (R^2^ = 0.8603, slope = 7.07 x 10^-5^) (**S3 and S4 Tables**). However, only 3 out of 8 *C. coli* and 5 out of 18 *C. jejuni* sub-lineages exhibited strong temporal signal with R^2^ >0.5.

Bayesian evolutionary analyses were performed on sub-lineages where temporal signal was strong (R^2^ > 0.5) using BEAST2 [45]. This excluded species-wide datasets but included 3 *C. coli* and 5 *C. jejuni* sub-lineages for which rate estimates were obtained (**S3 and S4 Tables**). Mean rate estimates were similar for all 3 *C. coli* lineages averaging at 7.82 x 10^-4^ s/s/y but varied from 8.20 x 10^-5^ (ST-661 clonal complex) to 1.00 x 10^-3^ s/s/y (ST-22 clonal complex) for *C. jejuni* (**S3 and S4 Tables**). In part because of the poor temporal signal in the species-wide analyses and most sub-lineages, we developed an alternative method using paired isolates.

### Sampling matched isolate pairs allows estimation of the rate of nucleotide change

Nucleotide change (NC) is introduced into the bacterial genome by recombination resulting from HGT, and point mutation. For clarification, we use the empirical term ‘nucleotide change’ to describe any nucleotide variation resulting from these two processes, consistent with previous studies [46]. Estimation of molecular clock rates requires comparison of isolates from related, or preferably the same, lineages that have accumulated NCs over time. To achieve this there is a necessary balance between maximizing the time between sampling and accumulated NCs whilst ensuring comparisons are made between related strains. Therefore, we plotted NC difference against time difference to determine criteria for choosing comparable isolate pairs (Fig 2). The sample time difference was chosen to maximize the time between sampling and the number of comparable pairs belonging to the same lineage. Pair selection criteria were standardised for both species so that isolate pairs were included where the sampling time difference was >8 years and there were <5000 SNPs between them (Fig 2**, S2 and S5 Tables**). Based upon these criteria, there were 18 *C. coli* and 74 *C. jejuni* isolate pairs (**S5 Table**). However, for consistency between the species we chose the 20 *C. jejuni* pairs with the highest nucleotide identity, hence those with the strongest evidence of recent common ancestry. Therefore 18 *C. coli* and 20 *C. jejuni* pairs comprised the dataset for NC rate calibration. These belonged to the ST-21, ST-22, ST-45, ST-1332, ST-828 clonal complexes and isolate pairs had a difference in sampling date of 8 to 11 years (*C. coli*) and 8 to 36 years (*C. jejuni*) (Fig 3**, S2 and S5 Tables**).

**Fig 2.**
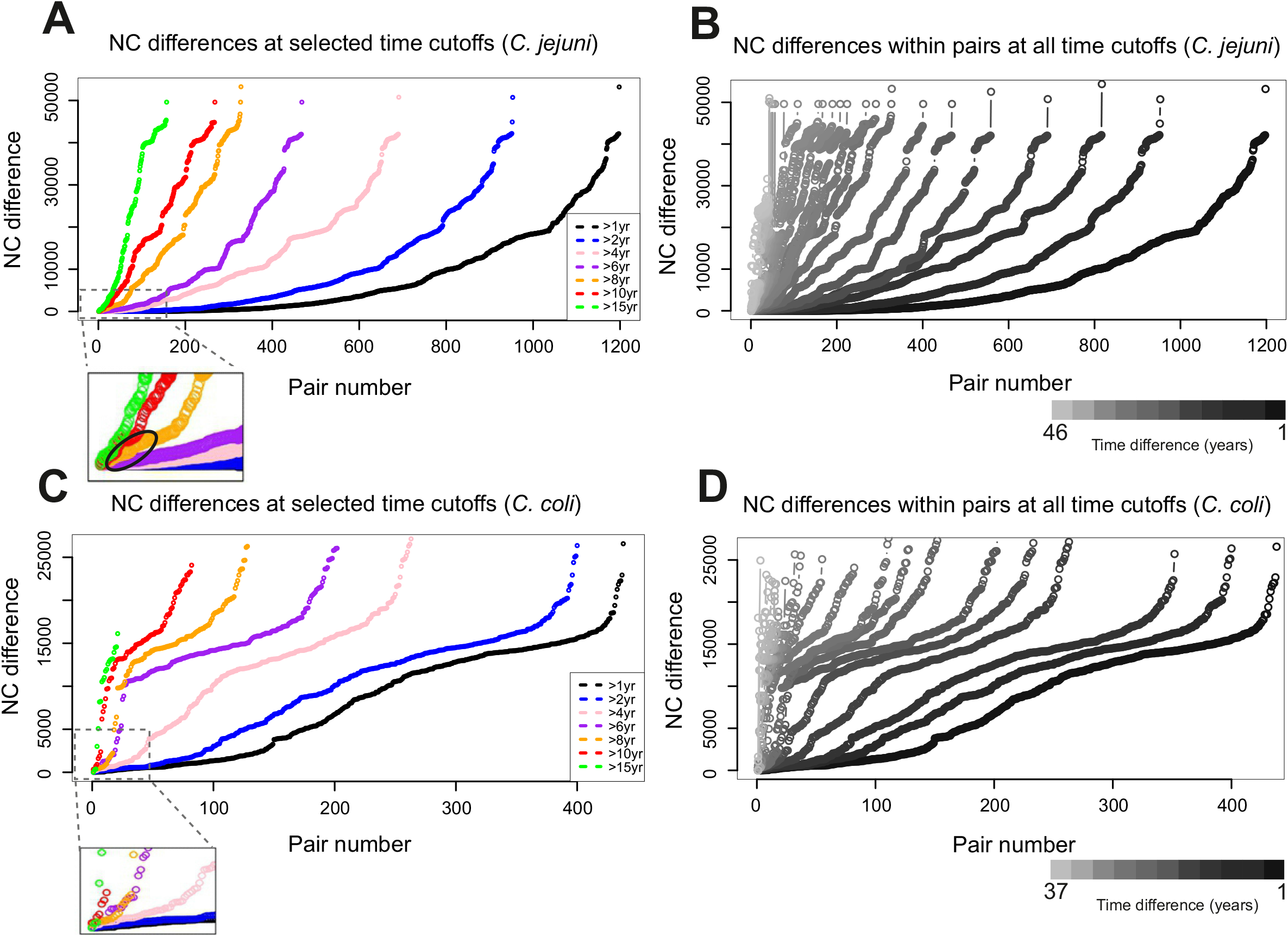
Pair selection criteria curves for inclusion in rate estimates. Visual representation of possible pairs of isolates at all time cut-offs across the sample time frame for *C. jejuni* (B) and *C. coli* (D). As time difference between pairs increases, distinguishing between individual curves becomes distorted. Therefore, a selection of years were plotted (A and C) (black = all pairs >1 year difference, blue = >2 years, pink = >4 years, purple = >6 years, orange = >8 years, red = >10 years, green = >15 years). All isolates were paired with the nearest isolate (genetic distance), matched according to difference in year of isolation (coloured lines) for both *C. jejuni* (A) and *C. coli* (C) (orange line). Dashed boxes (A and C) show magnified images of the closest pairs from all curves. The 20 and 18 pairs used in rate calibration are highlighted by a black oval (A) and every orange pair in dashed box (C). Grey scale bars (B and D) indicate the time difference cut-off of each curve for every time point in the sample date frame.

**Fig 3.**
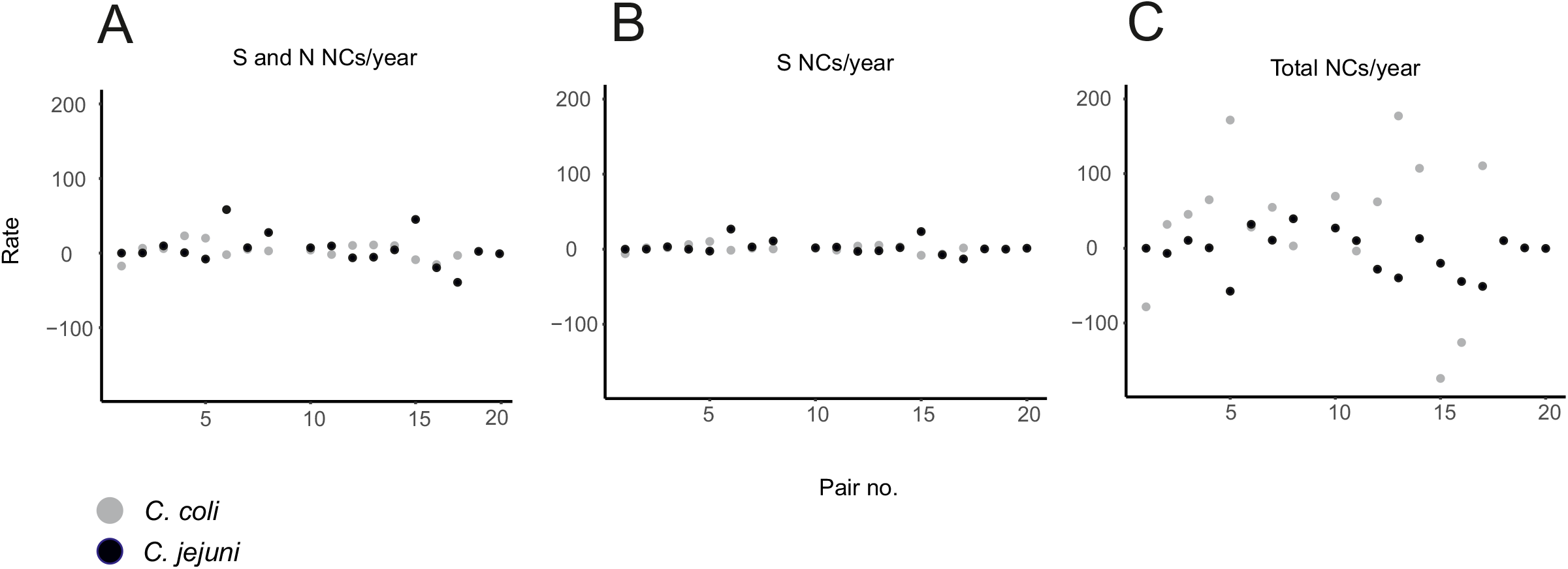
Scatter plots of individual pair NC rates for *C. coli* and *C. jejuni*. Three different NC rates were determined per pair for *C. coli* (grey circles) (18 pairs) and *C. jejuni* (black circles) (20 pairs): (A) all S (synonymous) and N (nonsynonymous) NCs, excluding those in recombined regions; (B) S NCs only, excluding those in recombined regions (molecular clock); (C) all NCs including those resulting from recombination (total NC rate). Variation was gratest among total NC rates. This demonstrates the impact of recombination, introducing the majority of NC’s into *Campylobacter* genomes. All remaining information on isolate pairs can be found in S2 Table.

Estimation of a molecular clock rate requires that NCs accumulate over time, defined here as NCs per site per year (s/s/y). It is also possible that branch shortening can occur where there are fewer NCs in the more recent isolate of a pair. While not specifically describing branch shortening, negative rates of NC have previously been observed [44]. In this study, 13 out 18 *C. coli* and 11 out of 20 *C. jejuni* isolate pairs exhibited branch lengthening, that is to say more total NCs (within and outside recombined regions) were found in the more recent isolate (**S2 and S6 Tables**). Only pairs having undergone measurable evolution (branch lengthening) were included in further analysis of the accumulation of NCs over time. While this may inflate our estimate of the NC rate, it was necessary to ensure a positive rate for calculating the number of effective lineages. For measurably evolving isolate pairs, the total NC rate was calculated as well as the rates within and outside of recombined regions (Table 1**, S7 Table**). The mean NC rate for non-recombined regions was 6.36 x 10^-6^ and 8.45 x 10^-6^ s/s/y but ranged from 1.60 x 10^-6^ – 1.50 x 10^-6^ and 1.00 x 10^-7^ – 3.60 x 10^-5^ s/s/y for *C. coli* and *C. jejuni* respectively, or 11.46 and 13.53 average NCs per genome per year (s/g/y) (Table 1).

**Table 1.** Average rate calibrations for *C. coli* and *C. jejuni*.

| Rates of nucleotide change | *C. coli* |  |  | *C. jejuni* |  |  | Units** |
| --- | --- | --- | --- | --- | --- | --- | --- |
|  | Min | Mean | Max | Min | Mean | Max |  |
| Total* | $1.7 \times 10^{-6}$ | $6.3 \times 10^{-5}$ | $3.0 \times 10^{-4}$ | $4.8 \times 10^{-8}$ | $8.8 \times 10^{-6}$ | $2.3 \times 10^{-5}$ | s/s/y |
| S and N (exc. rec) | $1.6 \times 10^{-6}$ | $6.4 \times 10^{-6}$ | $1.5 \times 10^{-6}$ | $1.0 \times 10^{-7}$ | $8.5 \times 10^{-6}$ | $3.6 \times 10^{-5}$ | s/s/y |
| S (inc. rec) | $4.9 \times 10^{-6}$ | $3.1 \times 10^{-4}$ | $1.8 \times 10^{-3}$ | $2.1 \times 10^{-7}$ | $1.9 \times 10^{-6}$ | $7.6 \times 10^{-6}$ | s/s/y |
| S (exc. rec) (molecular clock) | $2.1 \times 10^{-7}$ | $2.4 \times 10^{-6}$ | $7.7 \times 10^{-6}$ | $3.8 \times 10^{-8}$ | $3.4 \times 10^{-6}$ | $1.7 \times 10^{-5}$ | s/s/y |
| N (inc. rec) | $4.4 \times 10^{-8}$ | $2.4 \times 10^{-5}$ | $1.1 \times 10^{-4}$ | $5.0 \times 10^{-8}$ | $1.4 \times 10^{-6}$ | $4.8 \times 10^{-6}$ | s/s/y |
| N (exc. rec) | $4.8 \times 10^{-7}$ | $3.2 \times 10^{-6}$ | $7.8 \times 10^{-6}$ | $3.1 \times 10^{-7}$ | $4.8 \times 10^{-6}$ | $1.6 \times 10^{-5}$ | s/s/y |

\*all NCs from including (inc) and excluding (exc) recombination (rec); \*\*NCs per site per year (*C. jejuni* = 1.6 Mbp, *C. coli* = 1.8 Mbp); S = synonymous NCs, N = nonsynonymous NCs

### Recombination drives molecular evolution in *Campylobacter*

NCs in coding sequence based on gene definitions in the reference *C. coli* (YH501) and *C. jejuni* (NCTC 11168) isolate genomes introduced an average of 1569 and 242 NCs in *C. coli* and *C. jejuni* paired genome datasets respectively. Of these, an average of only 222 (*C. coli*) and 106 (*C. jejuni*) were inferred to be the result of point mutation, with the remainder resulting from recombination (**S2 Table**). Recombination is therefore the major source of sequence variation in both species (Fig 4**, S2 Table**), introducing nearly six times as many NCs in *C. coli* than in *C. jejuni –* consistent with previous estimates based upon MLST [47].

**Fig 4.**
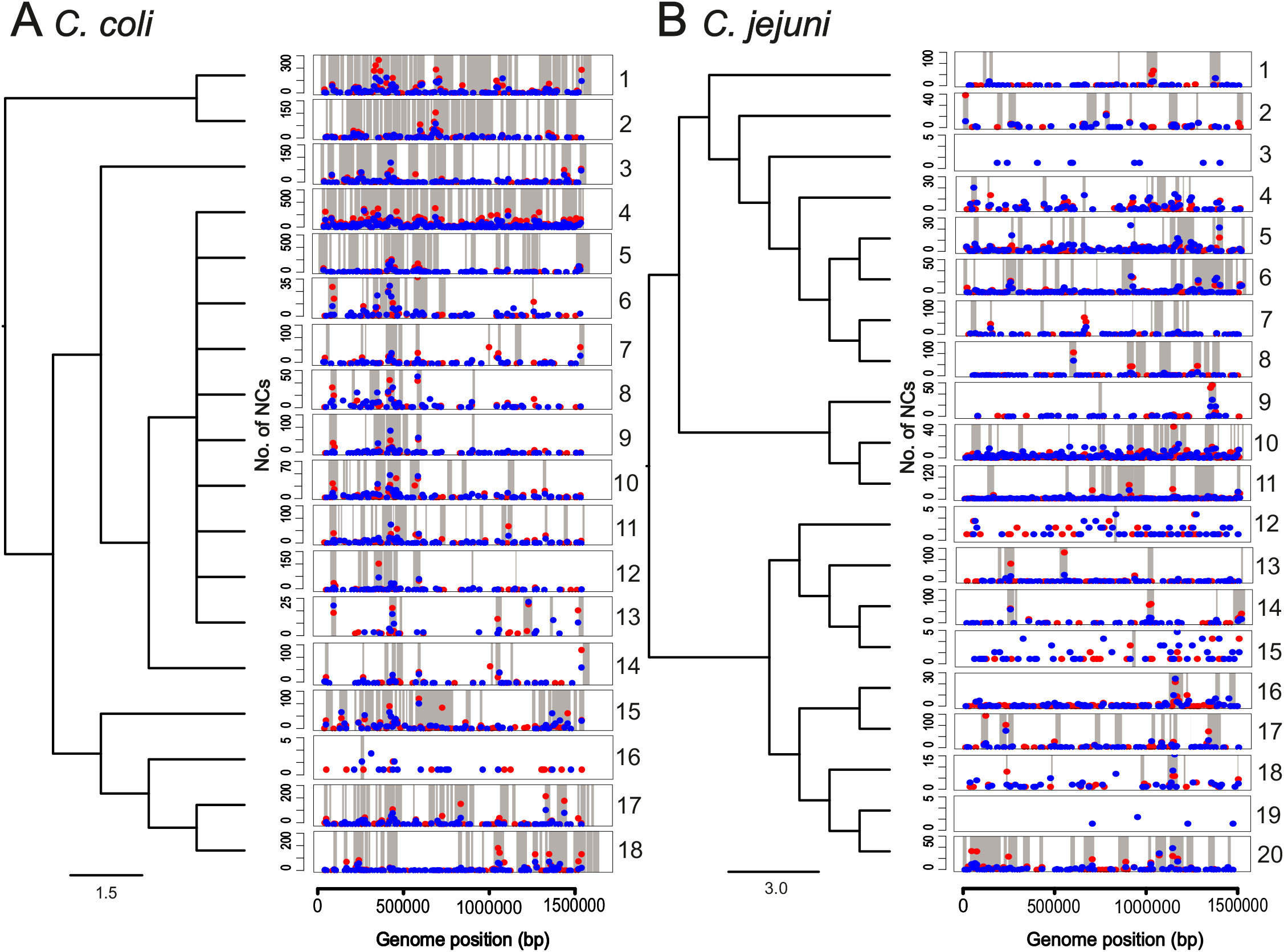
Mutation and recombination in *C. coli* and *C. jejuni*. Average genome-wide NC positions (red dots = synonymous NCs, blue dots = nonsynonymous NCs) per isolate pair in relation to inferred recombined regions (grey blocks). Each plot represents one pair of isolates considered in rate calibration for *C. coli* (A) and *C. jejuni* (B) and are ordered according to S2 Table. *y axis* = number of NCs in relation to particular bp position of the reference genome (*C. coli* = YH501, *C. jejuni* = NCTC11168) and varies between pairs. *x axis* = position of reference genome in bins of 10,000 bp. The cladogram shows the relatedness of isolate pairs based on nucleotide identity, scale bar indicates NCs per site. It is evident from both A and B that recombination is the main source of variation in *C. coli* and *C. jejuni*.

The effects of recombination on effective genotypes over successive generations were simulated for *C. coli* and *C. jejuni.* For both species, simulations provided results consistent with observations using our method on real data (Fig S5) in several ways. First, simulation of *Campylobacter* evolution under different recombination rates demonstrates how elevated nucleotide change (due to recombination) effects the number genotypes in successive generations. Second, the increase in the number of genotypes (and isolate pairs) is observed in both real and simulated data with evidence that rate slows over time. Third, the number of genotypes carried over to the next generation was higher for *C. coli* than *C. jejuni* in simulations at different recombination rates.

To assess the effect of NCs on amino acid sequences we quantified non-synonymous (N) and synonymous (S) NCs and determined the ratio per site (*dN/dS*) for all isolate pairs in recombined and non-recombined sequence (**S2 Table**). Point mutation on average accounted for an unequal amount of N and S polymorphism both within and between species (*C. coli*, N = 99, S = 123; *C. jejuni*, N = 63, S = 43) (**S2 Table**). While recombination introduced many more NCs than point mutation, in both species these were biased towards synonymous changes. Specifically, around six times as many S than N NCs were introduced by recombination in *C. coli* and approximately twice more in *C. jejuni* (*C. coli*, N = 546, S = 801; *C. jejuni*, N = 59, S = 77) (**S2 Table**). Overall, average *dN/dS* ratios were consistent between species within recombined (*C. coli* 0.492, *C. jejuni* 0.490) and non-recombined (*C. coli* 0.594, *C. jejuni* 0.509) portions of the genome. However, because of the relative importance of recombination (*r/m* = 37.240 (*C. coli*), *r/m* = 5.098 (*C. jejuni*)), on average N NCs were similar for *C. jejuni* from recombination and point mutation (59 and 63 respectively). However, recombination introduced 5.5 times more N NCs than point mutation in *C. coli* (**S2 Table**). Variation in *dN/dS* was observed between isolate pairs but was mostly indicative of purifying selection (*dN/dS*<1). Evidence of positive selection (*dN/dS*>1) was only observed within recombined sequence in 6 isolate pairs (**S2 Table**). It is important to note that, while *dN/dS* comparisons have been made between closely related bacteria within the same species [48, 49], this method was originally intended for between species comparisons [50].

Additional analysis of the distribution of recombination events revealed that an average of 13% (*C. coli*) and 2% (*C. jejuni*) of the genome has undergone recombination in at least one isolate pair since divergence from the common ancestor of each sub-tree (**S2 Table**). Recombination was distributed across the genome in both species but was elevated in certain regions of *C. coli* introducing more NCs at potential recombination hotspots [51]. However, recombination remained the main source of variation in both species (Fig 4).

### Molecular clock estimates for *C. coli* and *C. jejuni*

Molecular clock estimates require that NCs accumulate at a consistent rate over time. We maximized the chance of identifying this signal in several ways. First, genomic variation within recombined regions was discounted as multiple NCs can be introduced in a single evolutionary event – distorting clock estimates [39,47,52]. Second, non-synonymous NCs were discounted as selection may be more likely to influence the frequency of variation at these sites. Third, only pairs in which the most recently sampled isolate contained more NCs (branch lengthening) were used as they displayed measurable evolution over time. Based on these criteria, a similar average molecular clock rate was obtained for *C. coli*, 2.4 x 10^-6^ s/s/y (4.27 s/g/y), and *C. jejuni*, 3.4 x 10^-6^ s/s/y (5.42 s/g/y) (Table 1) but ranged from 2.1 x 10^-7^ –7.7 x 10^-6^ and 3.8 x 10^-8^ – 1.7 x 10^-5^ s/s/y.

### Coalescence and maintenance of lineages over time

Molecular clock estimates can vary within a population. Therefore the applicability of generalized clocks depend upon how much of the population has been sampled. To quantify this we estimated the average total NC rate (***µ***) (*C. coli* = 77.292 s/g/y, *C. jejuni* = 14.101 s/g/y), including all NCs within and outside recombined sequence. These rates were used to determine the number of coalescences in the population at a given time point (here referred to as ‘*effective lineages’*) within the dataset. The maximum timeframe for comparison was 37 years for *C. coli* and 46 years for *C. jejuni* (short in evolutionary terms). This provided information about the number of ancestral strains and the rate of turnover of lineages within the dataset. The total number of potential pairs without accounting for genetic similarity (***Y***), was equal to the square of the total number of isolates (*n*^2^) divided by two (to avoid double counting of isolate pairs), 180,600 and 1,663,488 for *C. coli* and *C. jejuni* respectively.

Having determined the total NC rate, we were able to predict the expected number of NCs over a given period of time. For example, 14 in 1 year for *C. jejuni.* We then subsampled all isolate pairs (***Y*)** to determine how many isolate pairs had ≤14 NCs between them – 76 isolate pairs. This is the possible number of isolate pairs that have arisen in 1 year. This process was repeated for each time cut-off, up to a maximum of 37 and 46 years for *C. coli* and *C. jejuni* respectively (**S8 Table**), to give the number of possible pairs for every time cut-off (***X***) (Figs 2B and 2D). Dividing ***Y/X*** resulted in the number of coalescences (*effective lineages*) at a given time interval in the past (***Z***) (**S8 Table**). For example, if the total NC rate was 14 s/g/y and we were interested in the number of birthdays within 5 years of our dataset, we would multiply the NC rate by 5 to result in 70 NCs of evolution over 5 years. The number of *potential pairs* (***Y*** = 1,663,488) / *possible pairs* (***X*** = 174) = ∼9,560 coalescences (ancestors) within this time period (**S4 Fig**, **S8 Table**).

The number of effective lineages at a given time-point can also be interpreted as the number of lineages that gave rise to those that are seen today. This provides valuable information about how the population is maintained over time and the extent to which it has diversified. For example, 1,263 *C. coli* lineages 37 years ago gave rise to an estimated 22,575 one year ago and 4,726 *C. jejuni* lineages 46 years ago gave rise to 21,888 lineages one year ago. This equates to an average increase in the number of effective lineages of 576 and 373 per year for *C. coli* and *C. jejuni* respectively. For *C. jejuni* it is clear that a considerable proportion (22%) of all lineages have been maintained throughout the 46 year sampling period and probably much longer (Fig 5). In contrast, only 6% of all effective lineages were present in the *C. coli* population 37 years ago. Perhaps the most striking finding is that the *C. coli* population has rapidly diversified in recent years. For example, there has been an 800% increase in the number of effective lineages in the last 10 years, over 3 times the rate of increase observed in *C. jejuni* (Fig 5**, S4 Fig**). This provides a basis for considering evolution in the wild but a longer sample time frame and more varied collection of isolates will improve representation of the natural *Campylobacter* population.

**Fig 5.**
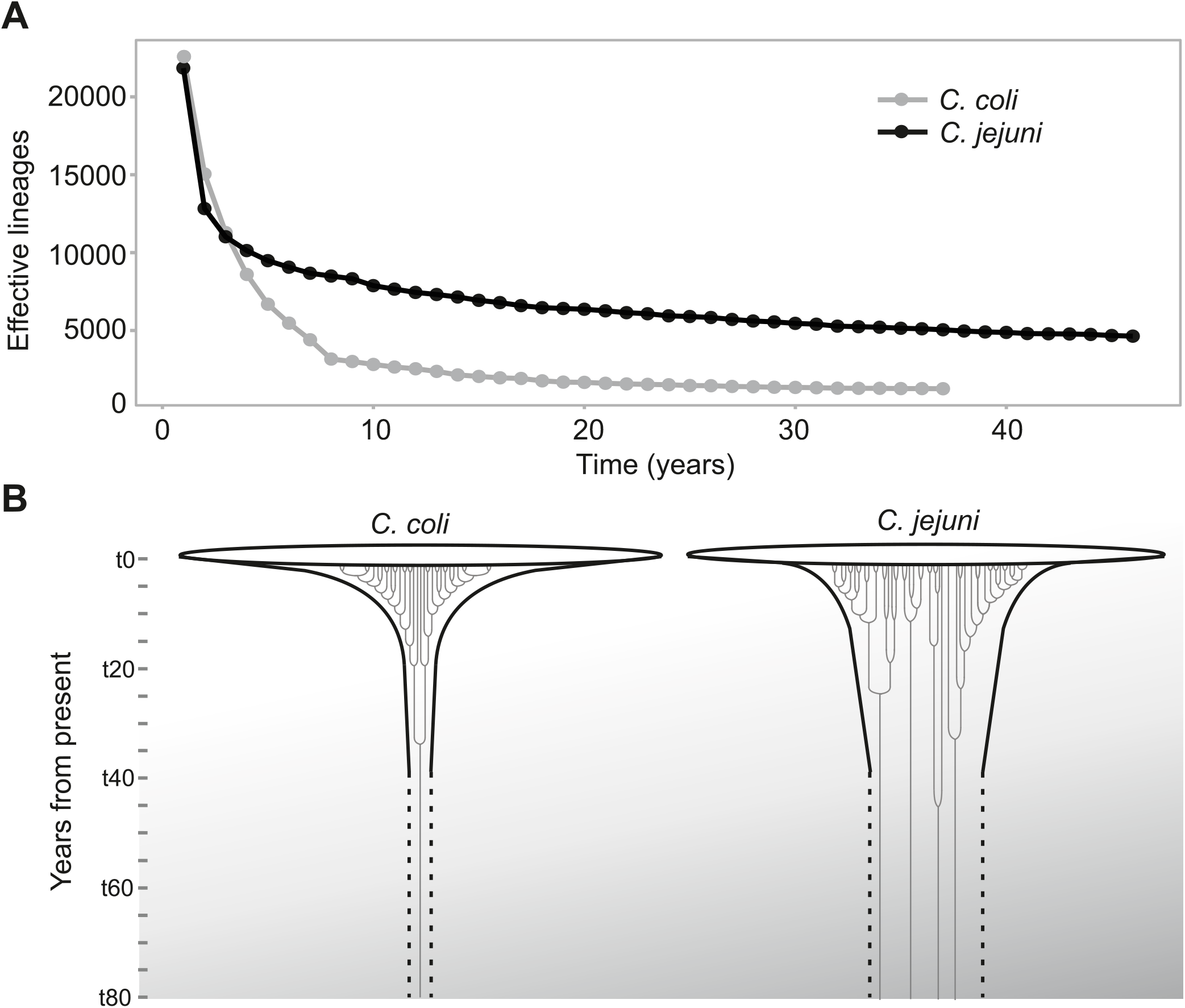
Lineage expansion in *C. jejuni* and *C. coli*. (A) Number of effective lineages (y axis) at each time point within the sample time frame (x axis) for *C. coli* (grey) and *C. jejuni* (black). (B) Diagrammatic representation of lineage expansion in *C. coli* and *C. jejuni* showing contrasting lineage diversification scenarios.

## Discussion

The increasing availability of large genome datasets has great potential for improving molecular clock estimates in bacteria. However, significant challenges remain. While it is clear that the frequency of NCs can vary between different species and strains [6,7,10,24,44,53,54], the extent to which nucleotide variation represents an intrinsic molecular clock is often less apparent. Biological factors such as generation time, population size and recombination rate, and ecological factors including cellular responses to habitat variation or stress and the strength of natural selection, influence the rate at which NCs accumulate in populations [55]. Therefore, obtaining a robust molecular clock estimate from natural bacterial populations requires an appropriate sample frame and careful consideration of the nature of observed sequence variation.

In cases where there is a clear temporal signal among isolates, it may be possible to obtain a robust molecular clock estimate by applying models to large genome datasets [24]. However, analysing all *C. coli* and *C. jejuni* genomes gave a weak temporal signal. This is likely related to the population structure and biology of these organisms that is in stark contrast to many obligate human pathogens [24]. Consistent with many other zoonotic or environmental bacteria, *Campylobacter* is a diverse genus with multiple lineages (STs and clonal complexes) inhabiting multiple hosts/niches. This required a more targeted approach to microevolutionary analysis consistent with that used to investigate transmission in similarly variable organisms [21].

Sub-sampling within the isolate collection, sampled over 46 years, identified closely related pairs of isolates with divergent sampling dates. Clearly, calibration of the molecular clock requires that NCs accumulate over time. This was not the case in all isolate pairs. In some cases, the most recently sampled isolate had accumulated fewer NCs than the comparator strain leading to a negative NC rate. A negative slope of the root-to-tip regression line has been interpreted as evidence for a lack of temporal signal or for a large dispersion of the rate of change [24, 44]. Furthermore other studies have described the time-dependency of molecular evolution [56, 57] and it can be the case that deleterious NCs in the older isolate have been purged leading to differences in long and short term molecular clock estimates [48, 58]. To simplify interpretation in our study, isolate pairs where the most recent isolate had not accumulated NCs were excluded from the analysis. While this might inflate the NC rate estimate, in organisms with complex ecology such as *Campylobacter*, it is also possible that closely related isolates occupy different sub-niches and experience different selection pressures even when sampled from the same host.

Returning to the birthday problem analogy, considering the number of isolate pairs (equivalent to people with the same birthday) obtained from the original genome dataset can provide clues about the extent of lineage diversity in the natural population. Using total NC rates, we were able to assess the nature of coalescence across the sample time frame for each species. The coalescence we refer to here is equivalent to the number of ancestral strains at a particular time point (effective lineages) in the natural environment from which contemporary strains emerged. Effective population size (Ne) is commonly used to reflect the number of individuals in a population that contribute to subsequent generations [59]. This has been used to investigate bacteria but contrasting approaches can provide different estimates depending on the method used [53, 60]. The idea of effective lineages, described in this study, is related to Ne but is more specific for organisms that reproduce clonally. Rather than typical Ne estimates for sexual populations, where the mating of two individuals is largely independent of what happened in previous generations, the number of effective lineages in a bacterial population reflects the number of distinct lineages that will survive and therefore contribute to future generations. This provides information on the genetic inertia of the population, i.e. the limitations for future evolutionary pathways based on the number of successful ancestors at a particular point in time.

These analyses highlighted the importance of appropriate sampling when calibrating NC rates and can help in determining the extent to which samples represent the population as a whole. Specifically, by considering the number of coalescences in a random population, we can look back through the sample time frame to estimate the number of effective lineages across a randomly sampled dataset. For example, suppose we would like to know if our contemporary isolates have a common ancestor in 1980. We know that a proportion of these ancestors gave rise to the diversity we see today but many lineages would go extinct and therefore not contribute [61]. Based on an average NC rate of 14 s/g/y for *C. jejuni*, there would be 560 NCs over 40 years total evolution between a strain pair. So, one can then ask how many pairs are close enough genetically for that to be the case. This gives an estimate of the effective number of ancestors in 1980 that gave rise to the contemporary dataset - equivalent to the number of birthdays.

Our estimate of effective lineages aims to better account for both unobserved lineages and lineages that co-exist over a long period of time. Both of these estimates are strongly affected by the sample frame and, while we have relatively large isolate collections from livestock and humans, the same cannot be said for the vast number of other potential host and environmental sources. Therefore, our estimate of effective lineages is more useful as an estimate of the number of ancestors that have given rise to the diversity of strains that we see today. For *Campylobacter*, it is clear that multiple lineages have persisted over a long period of time. This indicates that although the population is large, the strains are not turning over at a particularly fast rate. The absence of lineage replacement is inconsistent with some models of bacterial evolution that predict periodic population bottlenecks [62] but this can be explained in several ways. First, it is possible that the 37/46 year sampling period in this study is not sufficient time to out-compete a rival strain. Second, bacteria occupy different niches that are sustained so strains are not in direct competition. Third, the fitness differences among strains are not great enough for one lineage to out-compete another.

As well as the maintenance of multiple lineages, there is also evidence for variation in the number of effective lineages that contributed to successive generations between the two major pathogenic *Campylobacter* species. While, demographic inference and estimates of the number of generations using BEAST were similar for *C. coli* and *C. jejuni* (**S3 and S4 Tables**), the number of effective lineages was shown to be consistently higher for *C. jejuni* throughout much of the sample frame. Furthermore, there was evidence for a rapid increase in the number of *C. coli* lineages that began around 8 years ago (Fig 5). The reason for this is unclear. The average synonymous NC rate estimates were similar for *C. jejuni* and *C. coli*, 3.4 x 10^-6^ and 2.4 x 10^-6^ s/s/y respectively, equating to approximately 5.4 (*C. jejuni*) and 4.3 (*C. coli*) NCs per genome per year. This is somewhat lower than previous estimates for *C. jejuni* calculated from 7-locus MLST (2.79 x 10^-5^ s/s/y) [47] but is within the range of molecular clock estimates calculated from genomic variation for *Enterococcus faecium* (9.35 x 10^-6^ s/s/y) and *Y. pestis* (1.57 x 10^-8^ s/s/y) [44]. Although average synonymous NC rates were consistent with other estimates, rates ranged from 2.10 x 10^-7^ to 7.70 x 10^-6^ s/s/y in *C. coli* and 3.80 x 10^-8^ to 1.70 x 10^-5^ s/s/y for *C. jejuni.* This implies uncertainty around the estimate. However, rate heterogeneity among lineages is not uncommon in bacterial species [63] potentially reflecting differences in the evolution and ecology of different species and strains.

While the average NC rate was consistent for *C. coli* and *C. jejuni*, the relative number of NCs introduced by homologous recombination and mutation (*r/m*) differed markedly, with on average 37-fold (*C. coli*), compared to 5-fold (*C. jejuni*), greater impact on sequence variation. HGT is known to be an important driver of genome evolution in *Campylobacter* [47, 64] but these estimates are considerably higher than previous ones using 7-locus MLST [11]. Recombination introduced nearly twice as many synonymous than non-synonymous NCs, but even taking this into account, recombined sequence accounted for around 79% of all non-synonymous variation. This highlights the importance of HGT in rapidly evolving *Campylobacter* genomes and provides evidence that recombination may have been an important factor in the recent diversification of *C. coli* [26,65,66], potentially associated with an adaptive radiation [67, 68] linked to the colonization of agricultural niches [69]. However, this should be balanced against the evidence of purifying selection within recombined sequence (*dN/dS* = 0.492 for *C. coli* and 0.49 for *C. jejuni)* and the removal of non-synonymous NCs through negative selection [48].

Finally, throughout this study we have emphasized the importance of sampling so that measures of molecular evolution are obtained by comparing recent samples with a true ancestor. The uneven distribution of lineages within the population and the possibility that they differ in key evolutionary measures (*r/m* and *dN/dS*), means that our molecular clock estimate may not be applicable to all *Campylobacter* lineages [21,71–72]. Perhaps this is best illustrated by considering two host-specialist *C. jejuni* lineages, one associated with chickens and the other with cattle [8, 26]. There are 19 billion chickens on earth compared to 1.3 billion cattle [73] and *C. jejuni* colonizes up to 80% of chickens [74] with much lower rates in cattle. As the efficiency by which natural selection acts on sequence variation is related to effective population size [41], the rate of fixation and removal of NCs will be much faster in *C. jejuni* in chickens. Furthermore, chickens have a higher body temperature than cattle therefore the *C. jejuni* will grow faster, have a shorter generation time, and accumulate NCs at a higher rate [9]. From this simple example, which ignores many important factors (eg. subniche structure, host transition bottle necking, resident microbiome) it is clear molecular evolution can be influenced by population-scale forces down to the physiology of the individual cell. The approach employed in this study goes some way towards mitigating effects that confound generalized molecular clock estimates. Focussing on well-defined closely related isolate pairs inevitably reduces the number of comparisons from which the mean molecular clock rate is estimated. However, consideration of the distribution of effective lineages within the population is essential for identifying robust molecular clock estimates in environmental bacteria with complex multi-host ecology and massive effective population sizes.

## Materials and Methods

### Isolate sampling, genome sequencing and assembly

The accuracy of molecular clock estimates are improved by analysing large numbers of isolates sampled over long time periods. While very large isolate genome collections exist for *Campylobacter* (see below), many of these were sampled within the last 20 years. Therefore, to extend the sample time frame we assembled an isolate collection comprising 53 isolates sampled between 1978 and 1985 (12 *C. coli*, 41 *C. jejuni*) derived from multiple sources (human, duck, cattle, dog, turkey, wild bird and pig (**S1 Table**)). These samples were streaked onto mCCDA (PO0119A Oxoid Ltd, Basingstoke, UK) with CCDA Selective Supplement (SR0155E Oxoid Ltd, Basingstoke, UK) and incubated at 37°C for 48h in a microaerobic atmosphere (85% N_2_, 10% CO_2_, and 5% O_2_) using CampyGen Compact sachets (Thermo Fisher Scientific Oxoid Ltd, Basingstoke UK). Single colonies from each plate was then sub-cultured onto Mueller Hinton (MH) (CM0337 Oxoid Ltd, Basingstoke, UK) agar and grown for an additional 48h at 37°C and stored in 20% glycerol stocks at −80°C.

DNA was extracted using the QIAamp DNA Mini Kit (QIAGEN, Crawley, UK), according to manufacturer’s instructions. DNA was quantified using a Nanodrop spectrophotometer before sequencing on an Illumina MiSeq sequencer using the Nextera XT library preparation kits with standard protocols. Paired end libraries were sequenced using 2 × 300 bp 3rd generation reagent kits (Illumina). Short read data was assembled using the *de novo* assembly algorithm, SPAdes (version 3.10.0 35) [75] generating an average of 49 contigs (range: 2 −115) for a total average assembled genome size of 1.69 Mbp (range: 1.62-1.80). The average N50 was 189,430 bp (range: 81,283-974,529). These isolate genomes were augmented with 1,783 *C. jejuni* and 589 *C. coli* genomes archived in BIGSdb [76] representing isolates sampled from multiple sources (human, cattle, chicken, cat, dog, duck, environmental waters, farm environments, geese, lamb, rabbit, sand, seal, wild birds, turkey, pig) between 1970 and 2016 (**S1 Table).** The total isolate collection comprised 2,425 *Campylobacter* genomes, including *C. jejuni* belonging to 286 STs and 36 clonal complexes, and *C. coli* to 125 STs and 1 clonal complex. For *C. coli*, much of the genetic variation is within three ancestral clades, thought to have diversified before major introgressions with *C. jejuni* [66, 77–78]. However, isolates from these clades are not a major cause of human infection. In fact, ∼96% of all disease-causing *C. coli* strains (PubMLST, 16/08/2021), belong to the (introgressed) ST-828 clonal complex analysed in our study. For this reason we focused on the ST-828 complex [38]). All assembled genomes and raw reads have been deposited in the NCBI repository associated with BioProject: PRJNA524315. Individual accession numbers can be found in **S1 Table**. Assembled genomes of all isolates used in the study are available in FigShare DOI: 10.6084/m9.figshare.7886810.

### Simulating evolution in *Campylobacter* genomes

We performed forward-time simulations of neutral evolution on populations of *C. coli* and *C. jejuni* using a Wright-Fisher model in order to test the impact of recombination and point mutation on the effective genotypes over successive generations (time) using Bacmeta [79]. Starting population sizes of 1000 bacteria represented by 10 loci of length 1 kb were simulated for 60,000 generations to represent ten years of *Campylobacter* doubling time [80]. Mutation rates were set at 1 x 10^-5^ and 1 x 10^-6^ base^-1^ generation^-1^ for *C. coli* and for *C. jejuni* respectively and simulations were run three times with high (35.641), medium (13.058) and low (0.191) *r/m* values. Recombination events were exchanged as complete loci (**S5 Fig**).

### *C. coli* and *C. jejuni* phylogenies and assessing temporal signal and ‘clock-likeness’

Phylogenies were constructed for 601 *C. coli* and 1,824 *C. jejuni* isolates (**S1 Table,** Fig 1). Gene-by-gene alignments were produced with comparison to reference *C. coli* (YH501, accession number NZ_CP015528.1) and *C. jejuni* (NCTC11168, accession number NC_002163.1) isolate genomes, sampled in 2016 and 1977 respectively. These reference genomes, belonging to the ST-828 and ST-21 clonal complexes, were chosen as those most commonly used in *Campylobacter* comparative population genomics studies [81, 82]. For some bacteria the use of a lineage specific reference genome may increase the pool of core genes identified. However, we do not to take this approach for two reasons. First, all *C. coli* in this study belong to a single lineage, the ST-828 clonal complex, so there would be little benefit to creating separate reference pangenomes for already closely related isolates. Second, much of the advantage of creating lineage-specific pangenomes in *Campylobacter* is lost due to extensive recombination between divergent lineages [47]. For example, after comparison of all isolate genomes to the ST-21 complex NTCT11168 *C. jejuni* reference, the average number of core genome SNPs identified for non-ST-21 complex strains (255) was not significantly different compared to ST-21 complex strains (217) (T-test, Welch’s correction, p=0.683).

Homology to the reference genome (*C. coli* YH501; *C. jejuni* NCTC11168) was determined using MAFFT, with default parameters of minimum nucleotide identity of 70% over >50% of the gene and a BLAST-n word size of 20. Core genes (2,014 for *C. coli* and 1,668 for *C. jejuni*), shared by all isolates within a species were concatenated and used to construct Maximum likelihood (ML) trees using FastTree version 2.1.8 and the Generalised time-reversible (*GTR*) model of nucleotide evolution [83]. Isolates were analysed to test for a temporal signal of the accumulation of genetic variation over time (**S1 and S2 Figs**). This was carried out prior to NC rate analysis using a phylogeny of genetic distances and sampling dates, and root-to-tip regression implemented in the software TempEst v1.5.1 [84] with the best fitting root selected to maximise the coefficient, R^2^. Core genome phylogenies contained dated-tip isolates sampled between 1970 and 2016 for *C. coli* and *C. jejuni*.

Root-to-tip regression analyses to identify a temporal signal in bacteria rely on the removal of recombination as this is often the main source of genetic variation. Such analyses have been performed on small genome collections (15-189 isolates) of multiple bacterial species [44]. However, for *Campylobacter*, this would not be sufficient to represent diversity within the species as there are >40 known clonal complexes within *C. jejuni* alone. Using existing methodology, it was not possible to exclude recombining regions and perform root-to-tip regression on the entire *C. jejuni* (1824 isolates) and *C. coli* (601 isolates) genome collections. To account for this, we also assessed temporal signal in the whole datasets after removal of outlier genes that likely reflect recent recombination. Specifically, using a comparative gene-by-gene approach [85], we determined the number of alleles per locus in all genomes through comparison with the reference genomes (*C. coli* YH501*; C. jejuni* NCTC 11168). Three new concatenated genome alignments were constructed for each species after removal of 1%, 5% and 10% of the most variable loci (most/least alleles per locus) of respective core genome length of 1522 (1.11 Mbp), 1398 (1.02 Mbp), 1238 (0.89 Mbp) (*C. coli*) and 1268 (1.2 Mbp), 1164 (1.04 Mbp), 1034 (0.91 Mbp) (*C. jejuni*) core genes. Consistent with analyses of complete masked core genome alignments, TempEst analysis of reduced alignments (1%, 5%, 10%) also revealed poor temporal signal. Therefore, we conducted analyses on sub-lineages within *C. coli* and *C. jejuni*.

In order to identify temporal signal using TempEst, individual subsets of *Campylobacter* lineages were chosen from our datasets for *C. coli* (ST-825, ST-827, ST-828, ST-830, ST-872, ST-1090 and ST-1541 of the clonal complex, ST-828) and *C. jejuni* (ST-1325, ST-1034, ST-206, ST-21, ST-22, ST-257, ST-353, ST-354, ST-42, ST-433, ST-45, ST-464, ST-48, ST-52, ST-574, ST-61, ST-658 and ST-661 clonal complexes). Core genes were identified for each sub-lineage by comparison to *C. coli* (YH501) and *C. jejuni (*NCTC 11168) reference genomes using MAFFT (as described above). Individual ML phylogenies were constructed from core gene alignments using FastTree. Recombined regions were inferred using ClonalFrameML with basic model parameters [86] and removed using a cfml-mask script and replaced with gaps (https://github.com/kwongj/cfml-maskrc). Recombination-masked trees of individual lineages were subsequently tested for temporal signal using TempEst software v1.5.1.

### Bayesian evolutionary analysis

Analysis with BEAST2 v2.5.0 [45] was performed using a method previously described for *Campylobacter* [8]. SNP alignments were constructed from variable sites of the whole datasets of 601 (*C. coli*) and 1824 (*C. jejuni*) isolates and for the sub-lineages which exhibited the strongest temporal signal using snp-sites v2.5.1 [87]. A time-scale phylogeny was constructed using BEAST2 v2.5.0 [45] based on variable sites, using the GTR +G4 model of DNA substitution. The relaxed log-normal clock with Bayesian skyline model was used as previously described [8]. Input xml files were prepared using BEAUti2 v2.5.0 [45]. A prior on the clock rate was set as a log-normal distribution with a mean value of 1 x 10^-6^ mutations per site per year with a lower value of 1 x 10^-8^ and an upper value of 1 x 10^-3^. Markov chains were run for 50 million generations, sampled every 10,000 generations with the first 5,000,000 generations (10%) discarded as burn-in.

### Selection of closely related isolate pairs

An ideal dataset for rate analysis would include isolate pairs with divergent sampling dates, sufficient to measure NC rates over time, while remaining close enough (clustering on the tree) to share reliable recent common ancestry. Furthermore we required as many pairs as possible for confidence in average rates. In order to achieve this, pairwise nucleotide identity and year between isolation date matrices were constructed separately for 601 (*C. coli*) and 1,824 (*C. jejuni*) isolates. Using a bespoke R script (https://github.com/SionBayliss/CallandMolClock), the distribution of nucleotide identity was determined for isolate pairs within sequential isolation date categories of 1 year or more (1-37 for *C. coli*, 1-46 for *C. jejuni*) by comparing every isolate to all other isolates (Fig 2). In each analysis, isolates were used only once as the ancestral or derived strain. Large numbers of divergent isolate pairs can be identified from distant time points reflecting genome evolution over time (Fig 2). However, our analysis required inference of recent common ancestry (<5,000 SNPs) to define isolate pairs. Specifically, we needed closely related isolates with a large difference between sample dates. To provide a quantitative basis for selecting pairs we compared the nucleotide identity of each isolate pair at given sampling time difference thresholds. Isolates with sample dates >8 years apart (Fig 2) were chosen for calibrating a rate of nucleotide change because there was an enrichment of closely related pairs at this threshold. An arbitrary threshold of <5000 SNPs was selected for candidate isolate pairs to be included in the NC rate calibration analysis based on the pair selection curves. The final number of pairs used in the analysis included the 20 most closely match pairs for *C. coli* and *jejuni* (Fig 2A).

### Recombination and nucleotide change inference

The raw reads of genomes (**S1 Table**) of isolate pairs (**S2 Table**) were mapped to the complete reference genomes: *C. coli* YH501 (accession: NZ_CP015528.1) and *C. jejuni* NCTC 11168 (accession: NC_002163.1) using the BWA-MEM algorithm [88]. Variants were called using Freebayes v1.1.0-dirty [89] and NC effects predicted and annotated using SnpEff version 4.3 [90] (**S6 Table**). These tools were included in the haploid variant calling pipeline, ‘snippy’ v3.0 (https://github.com/tseemann/snippy). Core genome sub-tree alignments were constructed using snippy-core. NCs introduced by point mutation and recombination were inferred on the alignments using Gubbins v2.4.1 (default settings) [91] for each isolate pair (**S6 Table**). The snippy pipeline was used to identify synonymous and non-synonymous NCs within and outside of inferred recombinant regions [90]. *dN/dS* ratios were calculated for sites across the core genome using the synonymous/non-synonymous analysis program (SNAP) v2.1.1 based on the Nei and Gojobori 1986 method [92, 93] (www.hiv.lanl.gov). Rates were calculated by reconstructing the NCs on internal branches (inferred using Gubbins) leading to the MRCA shared by a pair of isolates. By quantifying point mutation and recombination and synonymous and nonsynonymous NCs per branch, we were able to infer different molecular evolution rate estimates based on the difference in isolation time between isolates pairs (**S4 Fig C**). These included (i) the total NC rate, used to calculate the number of effective lineages and (ii) the rate of accumulation of synonymous NCs occurring outside of recombinant regions, used to estimate the molecular clock. Hotspots of recombination occurring across multiple isolate pairs were observed.

### Estimating the number of coalescences at yearly intervals (Birthday problem)

To consider the extent to which a given sample set represented genetic diversity within the population we developed a pipeline that calculated the number of coalescences (*effective lineages*, **Z**) at yearly time intervals (Z1, Z2, Z3….Zn) within the datasets. This is described by the equation **Z** = **Y/X,** Where: **Y** = all potential isolate pairs (n^2^/2); **X** = the number of possible pairs for each time interval (*t1, t2,t3….tn*) that is less than the predicted number of NCs that have occurred over a given time interval (*µ(t(1-n)*); ***µ*** = rate of nucleotide change; ***t*** = time interval between sampling dates, 1-46 and 1-37 years for *C. jejuni* and *C. coli* respectively. The resultant **Z** value for each time period is the estimated number of effective lineages (Birthdays) at each time cut-off, equivalent to the number of lineages sharing a common ancestor at a particular time interval (**S4 Fig**).

## Acknowledgements

All high-performance computing was performed on MRC CLIMB, funded by the Medical Research Council (MR/L015080/1). This publication made use of the PubMLST website (http://pubmlst.org/) developed by Keith Jolley and Martin Maiden [38] and sited at the University of Oxford. The development of that website was funded by the Wellcome Trust.

## Author contributions

JKC, SKS and DF designed the study and wrote the paper with BP. JKC, BP performed genomic analysis with input from HT, EM, SCB, EJF and JC. MB provided historical collection of isolates that EB cultured isolates for sequencing. EM, MDH and BP sequenced and assembled genomes. All authors contributed and approved the final manuscript.

## Supporting Information

**S1 Table. Isolate list information**

**S2 Table. Effects of NC from point mutation and recombination on isolate pairs**

**S3 Table. TempEst root-tip regression and BEAST analysis estimates (*C. coli*)**

**S4 Table. TempEst root-to-tip regression and BEAST analysis estimates (*C. jejuni*)**

**S5 Table. List of all possible pairs >8 years apart and <5000 NCs in difference for both *C. jejuni* and *C. coli***

**S6 Table. All annotated NC effects**

**S7 Table. Individual rate of nucleotide change estimates per pair for *C. jejuni* and *C. coli* (SNPs/year)**

**S8 Table. *C. coli* and *C. jejuni* “birthday problem” data and estimates of coalescence across sample time frame**

**S1 Fig. Root-to-tip linear regression of *C. coli* implemented in the software, TempEst.** Root-to-tip genetic distance (y axis) is correlated with sampling time (x axis) for phylogenies of (A) 601 *C. coli* and (B) 8 sub-lineages of the ST-828 clonal complex. Only 3 out of 8 sub-lineages had strong temporal signal (R^2^ > 0.5).

**S2 Fig. Root-to-tip linear regression of *C. jejuni* implemented in the software, TempEst.** Root-to-tip genetic distance (y axis) is correlated with sampling time (x axis) for phylogenies of (A) 1824 *C. jejuni* and (B) 18 sub-lineages representing *C. jejuni* clonal complexes. Only 5 out of 18 sub-lineages had strong temporal signal (R^2^ > 0.5).

**S3 Fig. Root-to-tip linear regression of *C. coli* and *C. jejuni* using TempEst software, after removal of the most variable core genes.** Core gene alignments were constructed from 601 (*C. coli*) and 1824 (*C. jejuni*) isolates. The most and least 1, 5 and 10% of variable loci (alleles/loci) were removed and phylogenies were analysed with Tempest. Root-to-tip genetic distance (y axis) is correlated with sampling time (x axis) to reveal poor temporal signal (R^2^ < 0.5) in all six scenarios.

**S4 Fig. Methods for calculating the number of effective lineages within the population.**

(A) The total number of *C. jejuni* and *C. coli* isolates in the population and all potential pairwise comparisons between putative ancestral (black) and contemporary (white) strains to give the total number of potential isolate pairs, Y. (B) Isolate pair selection based on divergent sampling date (>8 years) and a nucleotide identity threshold <5000 SNPs. (C) Total rate of nucleotide change (*µ*) calculated for all chosen pairs. The rate of accumulation of all synonymous (S), nonsynonymous (N) NCs, within (rec) and outside (mut) of recombined regions, was estimated since the most recent common ancestor (MRCA, red circle). The difference in NCs between each pair was divided by the difference in isolation years to give *µ*. (D) The total NC rate was used to estimate the number of NCs that were to accumulate over a time period and the number of possible isolate pairs at given time intervals (t*1, t2,t3….tn*) for each species.

**S5 Fig. Simulated migration of *Campylobacter* genotypes over successive generations using Bacmeta software.** Plots show the increase in the number of genotypes (y axis) over 60,000 generations (x axis) representing ten years of *Campylobacter* doubling time for *C. coli* (**A**) and *C. jejuni* (**B**). Forward simulations were run with mutation rates reflecting estimates from this study and three different *r/m* values (high (35.641) (dotted line), medium (13.058) (dashed line), low (0.191) (block line)) to monitor the effects of recombination on genotype frequency from one generation to the next. Simulated data was compared to the number of possible isolate pairs with fewer SNPs than predicted based on the total NC rate (X) at a given time for *C. coli* and *C. jejuni* (red dashed line).

## Notes

https://www.ncbi.nlm.nih.gov/bioproject/PRJNA524315

https://doi.org/10.6084/m9.figshare.7886810

https://github.com/SionBayliss/CallandMolClock

### Competing Interest Statement

The authors have declared no competing interest.

