## Supplementary figures and images for "Quantifying bacterial evolution in the wild: a birthday problem for *Campylobacter* lineages"

### S1 Fig

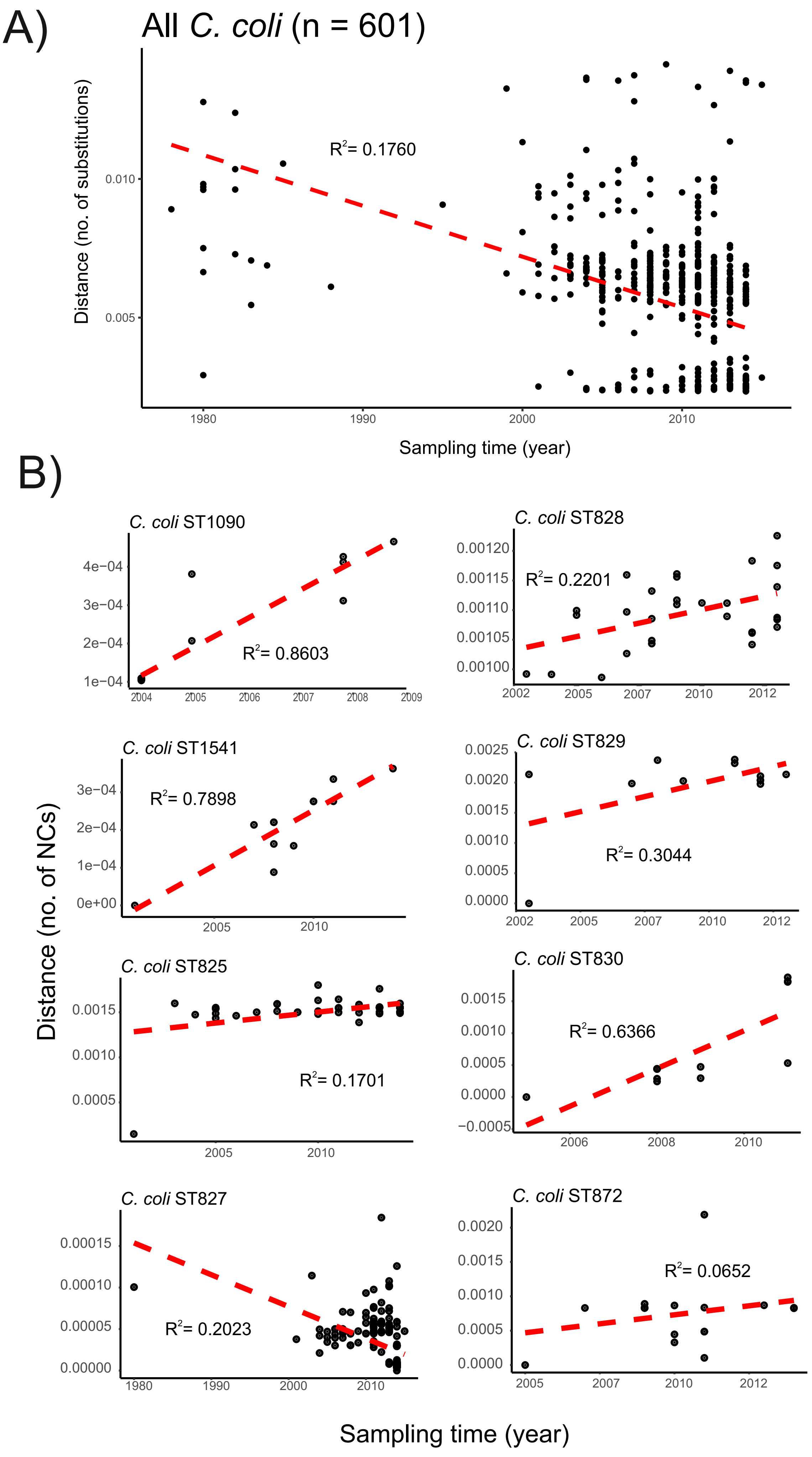

### S2 Fig

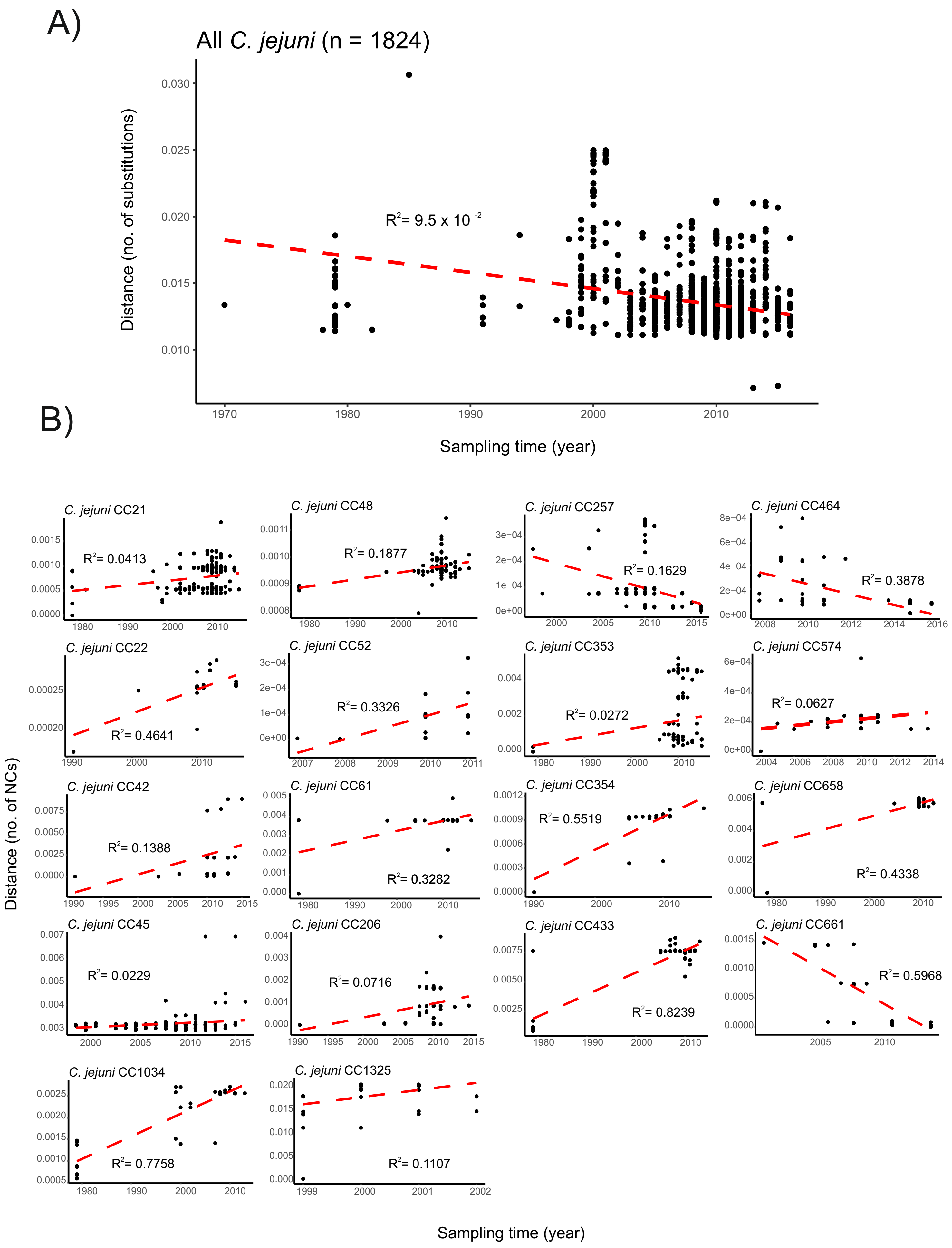

### S3 Fig

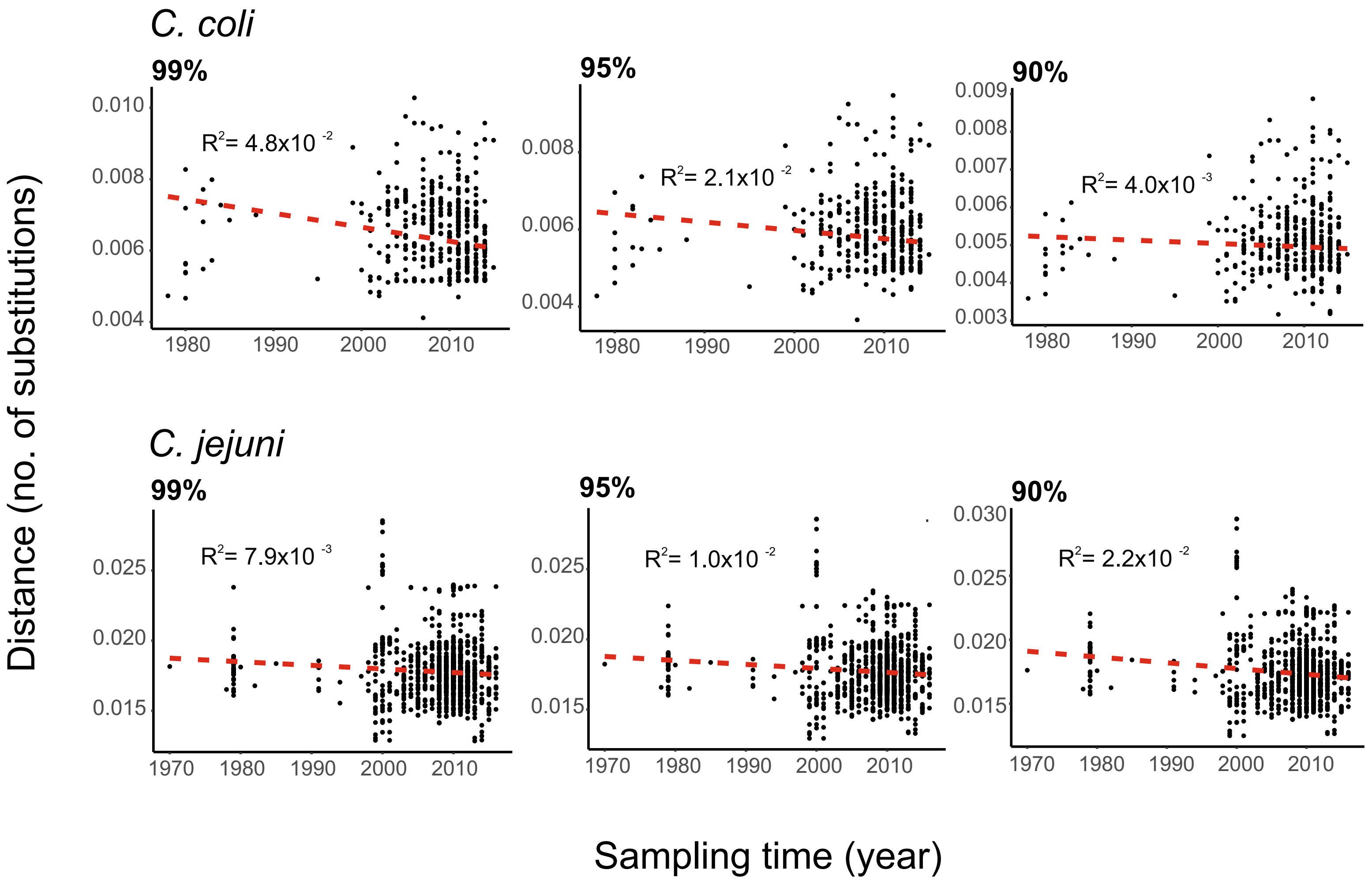

### S4 Fig

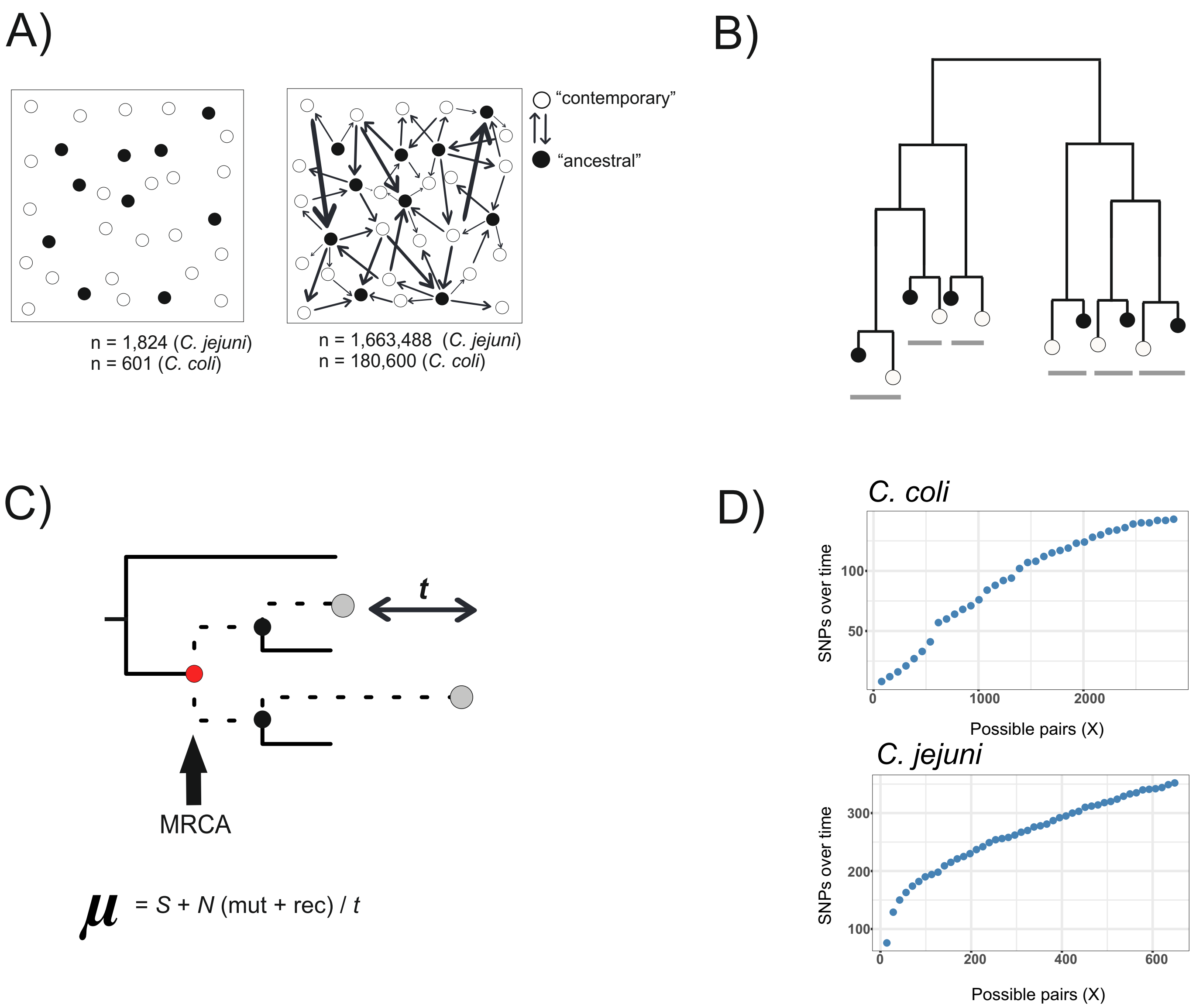

### S5 Fig

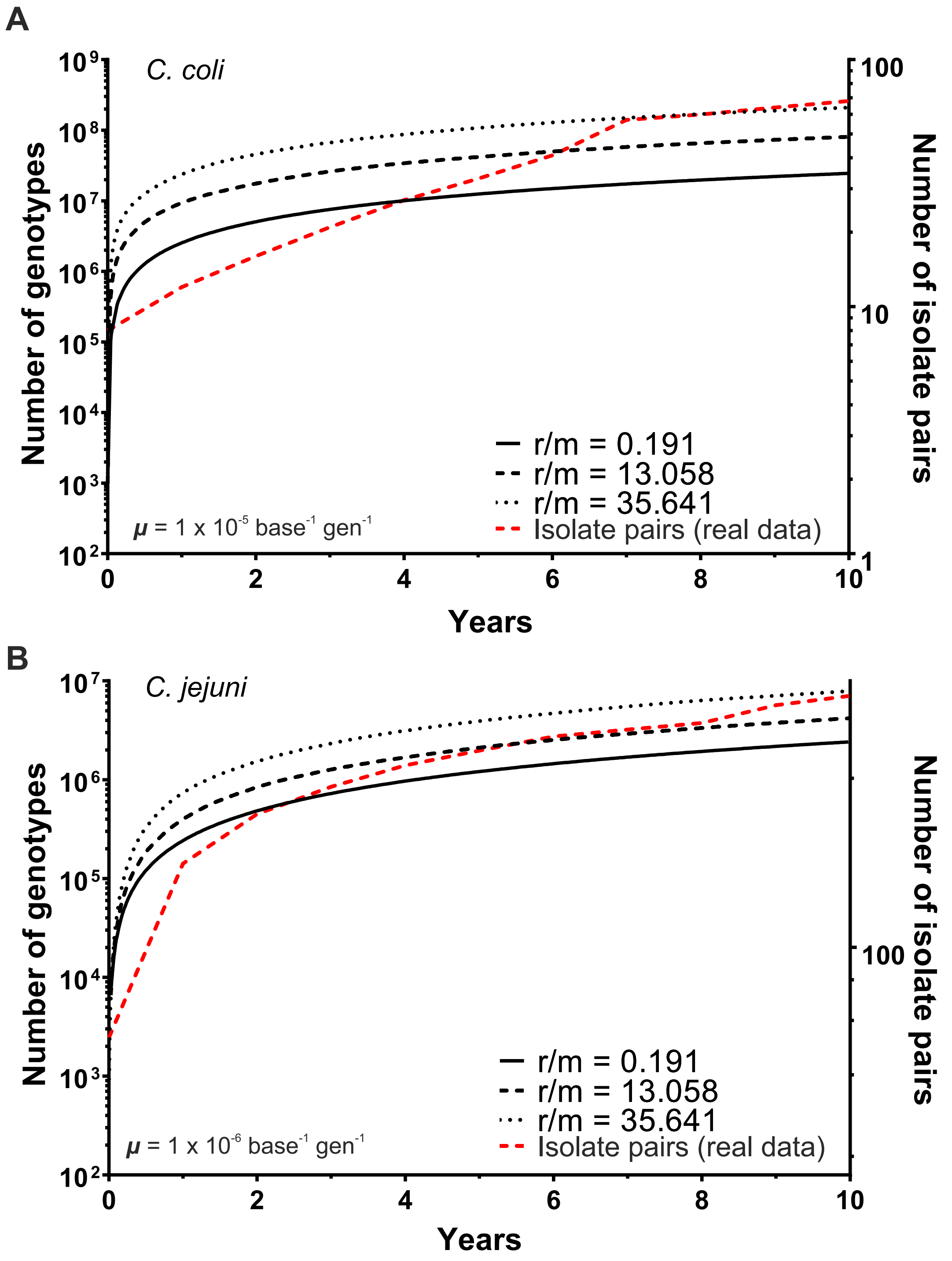
